# Relatively semi-conservative replication and a folded slippage model for simple sequence repeats

**DOI:** 10.1101/2020.02.28.970814

**Authors:** Hongxi Zhang, Douyue Li, Xiangyan Zhao, Saichao Pan, Xiaolong Wu, Shan Peng, Hanrou Huang, Ruixue Shi, Zhongyang Tan

**Author notes:** Co-first author (Hongxi Zhang, Douyue Li, Xiangyan Zhao, Saichao Pan). Corresponding author: Zhongyang Tan.

## Abstract

Simple sequence repeats (SSRs) are found ubiquitously in almost all genome, and their formation mechanism is ambiguous yet. Here, the SSRs were analyzed in 55 randomly selected segments of genomes from a fairly wide range of species, with introducing more open standard for extensively mining repeats. A high percentage of repeats were discovered in these segments, which is inconsistent with the current theory suggested that repeats tend to disappear over long-term evolution. Therefore, a mechanism is most probably responsible for continually producing repeats during replication to balance continuous repeat disappearance, which may makes the replicating process relatively semi-conservative. To improve the current straight-line slippage model, we proposed a folded slippage model involving the geometric space of nucleotides and hydrogen bond stability to explain the high-percent SSR occurrence, which can describe SSR expansion and contraction more reasonably. And analysis of external forces in the folding template strands suggested that the microsatellites tend to expand than contract. Our research may provide implements for contributions of microsatellites to genome evolution and complement semi-conservative replication.

## Introduction

Simple sequence repeats (SSRs), also referred as microsatellites, have attracted increasingly great interests in recent decades (Chen et al, 2010; Ellegren, 2004; Mandal et al, 2019; Morgante et al, 2002; Vinces et al, 2009a; Zhao et al, 2012), and have been widely analyzed in the genome sequences of eukaryotic prokaryotic and also viral genomes (Ellegren, 2004; Lin & Kussell, 2012; Morgante et al, 2002; Zhao et al, 2012). SSRs are the most variable genomic sequences, which tend to appear frequent variations in repeat-unit number instead of nucleotide substitution. And it may be a critical power accelerate the genomic evolution (Ellegren, 2004; Li et al, 2004), have roles associate with the host-adaptation and pathogenicity (Hood et al, 1996; Li et al, 2004), be relevant with the expression of genes and activity of promoters (Hannan, 2018; Vinces et al, 2009a), have relationship with many genetic diseases (Jain & Vale, 2017; Macdonald et al, 1993; Mirkin, 2007), and be observed with microsatellite instability (MSI) in many type of cancers (Bailey et al, 2018; Chan et al, 2019; Helleday et al, 2014; Kim et al, 2013).

Though SSRs have been comprehensively researched, there is actually no precise definition or wide-convinced standard for the extraction of SSRs all the time, which is usually based on setting the minimum numbers of the iterations for the mononucleotide to hexanucleotide SSRs based on empirical criterion (Chen et al, 2010; Ellegren, 2004; Li et al, 2004; Zhao et al, 2012). Majority of previous studies showed more interesting into the relatively longer repetitive sequences (Benson, 1999; Kelkar et al, 2010; Tian et al, 2011), and most studies usually used the threshold of 6, 3, 3, 3, 3, 3 for extracting mono- to hexanuleotide SSRs (Chen et al, 2011; George et al, 2012; Rajendrakumar et al, 2007; Zhao et al, 2011), while the very short repeat-motifs with smaller iterations were almost excluded, causing the neglect of their important significance (Fungtammasan et al, 2015; Hunt et al, 2016; Schmutz et al, 2014; Teh et al, 2017). In this work, the selected SSRs were extensively extracted with a wider extracting standard for extensive repeat-motif grabbing to investigate the essential occurrences of SSRs.

It is widely accepted that DNA slippage is thought to be the primary mechanism for driving microsatellites expansion or contraction, however, slippage involves DNA polymerase pausing, dissociation and re-association (Ellegren, 2004; Gadgil et al, 2016; Viguera et al, 2001), which may help to understand the expansion and contraction of long repeat sequences; it seems difficult to explain the remain of high percentage of short repeat sequences, and therefore, it is necessary to improve the slippage model more explicit to explain the generation of large amounts of short repeat sequences (Garcia-Diaz et al, 2006; Huang et al, 2017; Lai & Sun, 2003; Schlötterer & Tautz, 1992). It was suggested that the SSRs are most possibly born in the process of replication (Ellegren, 2004); replication is considered to be exactly semi-conservative with that the number of nucleotides in replicating chain is be precisely equal to that in template chain, and the replicating DNA molecule was shown as a straight molecule in vitro (Watson & Crick, 1953a; Watson & Crick, 1953b). Though it is well known that the DNA molecule is highly bent and packed in a super helix state within the nucleus, the replicating DNA molecule was also believed to be dragged to a straight molecule by the polymerase complex in vivo (Bell, 2011; Costa et al, 2011; Doublié et al, 1998; Kiefer et al, 1998). But there are a lot of environmental elements inside the nucleus which may disturb the polymerase complex, and these disturbances sometimes may affect the dragged straight DNA molecule returning to some extent of bent. The bent replicating DNA molecule is possibly related to the polymerase slippage for the occurrence of short SSRs. Here, we calculated the bent replicating DNA molecule with strictly considering the geometric space, the relationship between the phosphodiester bond and hydrogen bond, and also the stability of paired nucleotides; and proposed a folded replication slippage model for explaining repeats occurrence, which seems more reasonable to explain the remaining of high percentage short repeats in genomes, and also to explain the frequent microsatellite expansion and contraction. This work may also put forward some constructive suggestions for complementing the theory of semi-conservative replication.

## Results and discussion

### Genomes tend to produce short repeats

We analyzed 55 randomly-selected reported segment sequences covering from animal, plant, fungus, protist, bacteria, archaea and viruses (Table S1). The SSRs were extracted from all these segment sequences by using a threshold with minimum length of 3 base pairs or nucleotides. Though 2 iteration of di-, tri-, tetra-, penta- and hexa-nucleotide repeat sequence are usually ignored in most previous studies (Ellegren, 2004; Hunt et al, 2016; Schmutz et al, 2014; Teh et al, 2017; Zhao et al, 2012), we found they occurred in a very large number. It is difficult to consider them just as random sequences but not repetitive sequences, and it is also inappropriate to consider the iteration of 3 to 5 of mononucleotide repeats just as random sequences. Therefore, the threshold was set at 3, 2, 2, 2, 2, 2 in this study for exploring more comprehensive occurrence of SSRs, which could grab shorter simple repeats that never analyzed before, and another two thresholds were used to analyze these sequences for comparison. To test whether the SSRs under this threshold are random, we generated 55 mimic sequences with same size and nucleotide composition to the corresponding 55 reported sequences.

The analyzed data showed that the reported segment sequences are averagely 44.4% constituted, with SSRs, ranging from 36.4% to 60.0% under the new threshold (Fig 1A, Table S1). And comparing analysis also show the SSR content of these segments with average of 18.8% and 5.0%. These results indicate that all these segments remained high content of SSRs, because all these segments are randomly selected from their genomes, suggesting that the remaining high content of short SSRs is a general feature of all organism genomes after long time evolution, and also suggesting that few formerly well-studied repeats may only stand for the proverbial tip of the iceberg (Chen et al, 2010; Ellegren, 2004; Lin & Kussell, 2012; Morgante et al, 2002; Zhao et al, 2012). The null hypothesis test demonstrated that the percentages of SSRs in the generated segments are all lower than those in the reported segments, indicating that the high percentages of short SSRs are not randomly remained in all reported segments.

**Figure 1.**
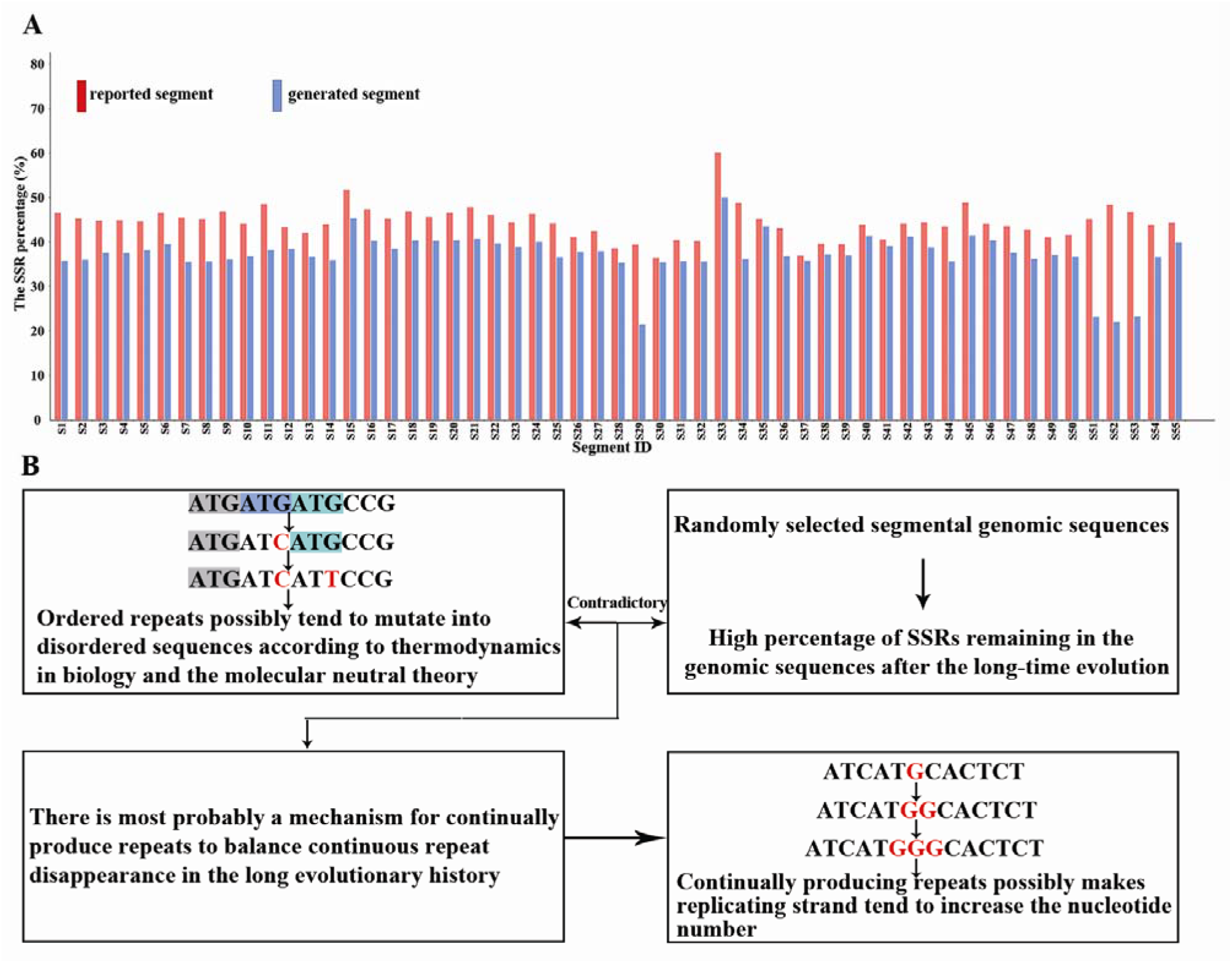
A high percentage of SSRs in genomes and genomes probably tend to produce repeats. A. SSR percentages of 55 randomly-selected reported segments and the control group, which consisted of the generated segments according to the sizes and nucleotide compositions of corresponding reported segments. B. Contradiction analysis of disappearance and high percentage of SSRs in the genomes.

Though the evolutionary mechanism of nucleotide sequences is still hotly debated by evolutionist, it is widely accepted that the genomic sequences are continually mutating forever; and the neutral molecular evolution and molecular clock theory suggested that the nucleotide substitution is constant over the evolution time; the thermodynamics in biology states that an isolated system will always tend to disorder (Bharadwaj et al, 2006; Kimura, 1977; Kimura, 1979; Margoliash, 1963; Zuckerkandl & Pauling, 1962; Zuckerkandl & Pauling, 1965). As the microsatellites are indeed ordered sequences, according to the former stated theories, the ordered repeats possibly tend to mutate into disordered sequences in the long evolutionary history without any selective pressure. Therefore, the repeat sequences should tend to disappear in genomes in the long evolution history. However, the remaining high percentage of SSRs in genomes is contradicted with the ideas of repetitive sequences tend to become no repetitive sequences. Thus, it can be inferred that there is most probably a mechanism for continually produce repeats to balance continuous repeat disappearance, and be responsible for the remaining of high percentage of short repeat sequences in genomes (Fig 1B).

Furthermore, the SSRs of small iteration numbers were observed to occur largely more than those of large iteration numbers in all analyzed segments (Table 1, Table S2), and this observation indicated that the SSRs of small iteration numbers maybe the basis for forming the SSRs of large iteration numbers, otherwise, it should be that the SSRs of large iteration numbers possibly are remained in higher percent level than or at least almost same level to the SSRs of small iteration numbers. Some of the longer SSRs also possibly mutate into short SSRs by contraction and point mutation as debated by many evolutionists (Ellegren, 2004; Kelkar et al, 2011; Mirkin, 2007), and these debates are possible because of that most of short repeats were not considered in their statistics; our observations generally suggested that most of the longer SSRs possibly evolved from short SSRs by expansion. So, the genomes possibly tend to produce short repeats by a continual repeat producing mechanism with the possibility of expansion a little more than that of contraction.

**Table 1.** The lengths (bp) of SSRs with different repeat unit types and different iterations in the segment of the reported human reference X chromosomal sequence at the location of 144822-231384 bp.

| Iteration | Mono<sup>a</sup> | Di | Tri | Tetra | Penta | Hexa | Total |
| --- | --- | --- | --- | --- | --- | --- | --- |
| I<sub>2</sub> | (18128)<sup>b</sup> | 10040 | 3540 | 2056 | 1250 | 480 | 17366 |
| I<sub>3</sub> | 9702 | 1782 | 288 | 156 | 45 | 18 | 11991 |
| I<sub>4</sub> | 3844 | 368 | 12 | 112 | - | - | 4336 |
| I<sub>5</sub> | 2095 | 120 | 15 | 20 | - | - | 2250 |
| I<sub>6</sub> | 600 | 24 | 18 | 0 | - | - | 642 |
| I<sub>7</sub> | 182 | 14 | -<sup>c</sup> | 28 | - | - | 224 |
| I<sub>8</sub> | 128 | 16 | - | 0 | - | - | 144 |
| I<sub>9</sub> | 54 | 18 | - | 36 | - | - | 108 |
| I<sub>10</sub> | 50 | 0 | - | - | - | - | 50 |
| I<sub>11</sub> | 55 | 22 | - | - | - | - | 77 |
| I<sub>12</sub> | 24 | - | - | - | - | - | 24 |
| I<sub>13</sub> | 65 | - | - | - | - | - | 65 |
| I<sub>14</sub> | 56 | - | - | - | - | - | 56 |
| I<sub>15</sub> | 45 | - | - | - | - | - | 45 |
| I<sub>16</sub> | 64 | - | - | - | - | - | 64 |
| I<sub>17</sub> | 0 | - | - | - | - | - | 0 |
| I<sub>18</sub> | 36 | - | - | - | - | - | 36 |
| I<sub>19</sub> | 19 | - | - | - | - | - | 19 |
| I<sub>20</sub> | 0 | - | - | - | - | - | 0 |
| I<sub>21</sub> | 42 | - | - | - | - | - | 42 |
| I<sub>22</sub> | 0 | - | - | - | - | - | 0 |
| I<sub>23</sub> | 23 | - | - | - | - | - | 23 |
| I<sub>24</sub> | 0 | - | - | - | - | - | 0 |
| I<sub>25</sub> | 25 | - | - | - | - | - | 25 |
| I<sub>26</sub> | - | - | - | - | - | - | - |
| I<sub>27</sub> | - | - | - | - | - | - | - |
| I<sub>28</sub> | - | - | - | - | - | - | - |
| Sum | 17109 | 12404 | 3873 | 2408 | 1295 | 498 | 37587 |

<sup>a</sup> Mononucleotide repeat (Mono), Dinucleotide repeat (Di), Trinucleotide repeat (Tri), Tetranucleotide repeat (Tetra), Pentanucleotide repeat (Penta), Hexanucleotide repeat (Hexa).
<sup>b</sup> The length of mononucleotide repeats with iterations of 2 was not included in this statistics and just used as the reference here.
<sup>c</sup> Beyond the largest iteration of this repeat unit type in corresponding analyzed segments were expressed as “-“.

### Relatively semi-conservative replication

It is well known that each base pair of DNA is one-to-one correspondence without other extra residue during replication in the double-helix model (Watson & Crick, 1953a; Watson & Crick, 1953b). And Meselson and Stahl have verified the replication of DNA chains is semi-conservative by the sedimentation techniques based on the diversity differential of DNA with different isotopes, also implicating that the number of nucleotides in replicating strand is consistent with that in template strand while processing complete replication (Meselson & Stahl, 1958). However, if the remained high percentage of short repeats is produced during replication process as described above, it certainly makes the base numbers of replication strand to be unequal to those of template strand, with one or several nucleotides/motifs being repeated and more than that in template strand. In vitro experiments also revealed the presence of repeats during DNA replication, and the nascent replication chain has a base increase (Doublié et al, 1998; Fungtammasan et al, 2015; Fungtammasan et al, 2016; Kiefer et al, 1998). And in this case, the replication process is possibly relatively semi-conservative and could be described as the following formula:

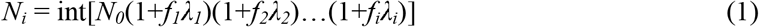

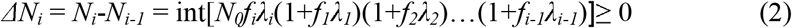

*N_0_*: The number of nucleotides in the initial template strand;

*N_1_*: The number of nucleotides in the replicating strand during No. *i* round replication;

int[]: Round the value to the lower integer;

*ΔN_i_*: The difference for the number of nucleotides between *N_i_*, and *N_i-1_*;

*λ_i_* (*λ_i_*,→ 0): The coefficient of occurring repeats during No. *i* round replication, and is most probably an infinitesimal with relating to the possibility of repeat sequence occurrence;

*f_i_* (0≤*f_i_*≤ 1): The fixation coefficient of repeat sequences during No. *i* round replication.

In general, the number of nucleotides in replicating strand is usually detected to be exactly equal to that in template strand, which is possibly because of the observed template strand being too short, for example, the total number of nucleotides in the initial template strand for stable PCR is up to two to three thousand nucleotides, in this case, we suppose *N*_0_ = 3000, *λ_1_* = 10^-5^, *f_1_* = 1, then the value of *ΔN_1_* will be 0 according to the formula (2), and therefore, *N_1_* = *N_0_*, causing the replicating strand to be no longer (or no shorter) than template strand, and the discovery of new-born repeat is unavailable; however, when the observed strand is long enough, then *ΔN_i_* is able to be larger than 1 at least, and it can be found that the number of nucleotides in replicating strand is different from that in template strand, for instance, we suppose *N*_0_ = 10^6^, *λ*_1_ =10^-5^, *f_1_*=1, then the value of *ΔN_1_* will be 10, in this case, the replicating strands probably have 10 nucleotides (or repeat-motifs) more than template strand do after this replication. Thus, the increased number of nucleotides may represent newly occurred repeat sequences.

The occurrence of SSRs will possibly encounter selective pressure, though it may be different in coding or non-coding regions, then, we use *f_i_* representing the fixation possibility of the newly born repeats facing with the selective pressure. The *f_i_* = 0 when the occurrences of new repeats are the lethal mutations and unable fixation in the organism, or may be excluded by DNA repair system (Jeggo et al, 2015; Mandal et al, 2019). The fixation coefficient is 0<*f_i_*<1 when the new SSRs are the deleterious but fixed in the genome within alive individuals, like Huntington’s disease (Macdonald et al, 1993). While the occurrences of new SSRs are the neutral mutations, the fixation coefficient should be 0≤*f_i_*≤1, and they are fixed or excluded depending on genetic drift. And the *f_i_* of beneficial mutations is 1, representing that the new SSRs may help the organism surviving. Therefore, the remaining high percentage of short repeats suggests that the replicating process possibly produce short repeat sequences frequently which may be fixed neutrally, beneficially, or deleteriously with diseases, and also suggests that the replication may be relatively semi-conservative.

### Folded slippage model

The nucleotide chains of various species tend to produce simple repeats during replication, and thus cause the number of the nucleotides in replication strand possible to be different from template strand after replication as discussed above. Moreover, how did simple repeats actually originate from is still a key argument topic (Ellegren, 2004; Kelkar et al, 2011; Torresen et al, 2019). The widely accepted mechanism of occurring SSRs is the replication slippage model, which is possibly easy to explain the expansion and contraction of longer SSRs, but possibly difficult to explain the much amounts of short repeats expansion and contraction. And the current slippage model is indeed a straight template strand model, without considering that the space is required for nucleotide base and also phosphodiester bonds are much stronger than hydrogen bond (Fig 2A) (Gao et al, 2004; Heyrovska, 2006), and also without considering what is the force to drive the replicate strand slippage. The straight replication slippage model has not given any clear suggestion, and it suggests that the SSRs possibly occurred by slippage occasionally (Gemayel et al, 2010; Leclercq et al, 2010; Mirkin, 2007; Ohshima & Wells, 1997). Actually, there are about 33 atoms in a nucleotide (A: 33, T: 33, G: 34, C: 31) (Alberts et al, 2002), and of course the nucleotide base need a certain space in nature. According to previous reports, we simplified a nucleotide space into an intuitive plane model, whose length is about 0.489 nm (length = (distance between the double helix 1.08 - Hydrogen bond length 0.102) / 2), and with a width of 0.34 nm which is the distance between each pair of bases (Fig 2A) (Gao et al, 2004; Heyrovska, 2006; Wang, 1993). We reconstructed the linear replication slippage model with a CAD geometric calculation by considering the space of bases (Fig 2B, Fig S1); if the slippage bubble has enough geometric space to accommodate the repeat bases, the phosphodiester bond should be elongated far more than 0.34 nm, while the phosphodiester bonds in DNA is actually much stronger than hydrogen bond (Fig 2A)(Wang, 1993). So it is impossible to form a slippage bubble by a larger elongation of the phosphodiester bonds for accommodating the repeat bases. Therefore, the straight slippage model is very difficult to the occurrence of short repeats, and it is most possibly necessary to improve the slippage model.

**Figure 2.**
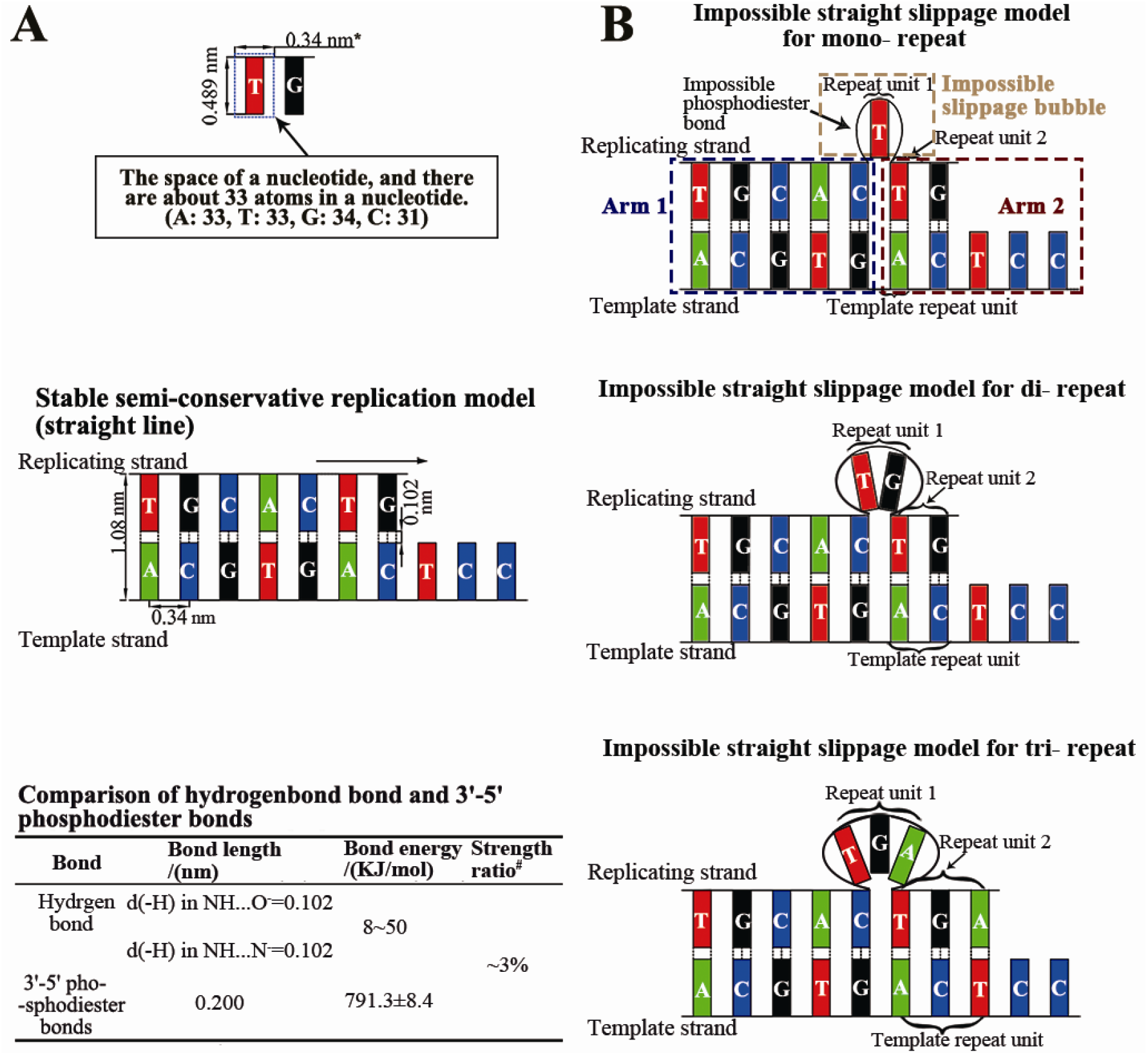
Straight strand models of semi-conservative replication and slippage. A. The space of a nucleotides was drawn. * indicates that those number is the theoretical values (top); The stable straight model of semi-conservative replication (middle); The comparison of hydrogen bond and 3’-5’ phosphodiester bonds (bottom)(Gao et al, 2004; Heyrovska, 2006; Wang, 1993). # indicates the strength ratio was calculated by the strength of hydrogen bond dividing that of phosphodiester bond. B. The impossible straight slippage models of mononucleotide, dinucleotide and trinucleotide repeats according to the strict geometric calculation of the space of a nucleotide and the stability of hydrogen and phosphodiester bonds.

Actually there is a fact which is widely ignored in replication slippage studies. The template strands are thought to be straight in all replication models, though it is the truth in general condition. It is also well known that the genomic DNA chains are very long and the space is too narrow in the nucleus (Fig 3A); for example, the total length of human genome is about 2 m (2×10^9^ nm), while the diameter of nucleus is beneath 10^5^ nm in human cell (Alberts et al, 2002); therefore, the genomic DNA chains are generally highly curved and folded in the nucleus as widely accepted. Indeed, the replicating molecule is believed to be a straight molecule (Bell, 2011; Costa et al, 2011; Doublié et al, 1998; Kiefer et al, 1998), and the replicating enzyme complexes usually straighten the template strand to be straight making the replicating strand well paired to finish the semi-conservative replication process (Costantino et al, 2014; Fragkos et al, 2015; Kiefer et al, 1998). However, there are a lot of environmental factors like temperature, viral proteins or diseases etc., which may disturb the normal works of the enzyme complexes. So, when the replicating enzyme complexes are disturbed by environmental factors, the replicating part DNA molecule may recover to some extent of curved or folded state, and then the template strand may also be some extent of curved or folded state.

**Figure 3.**
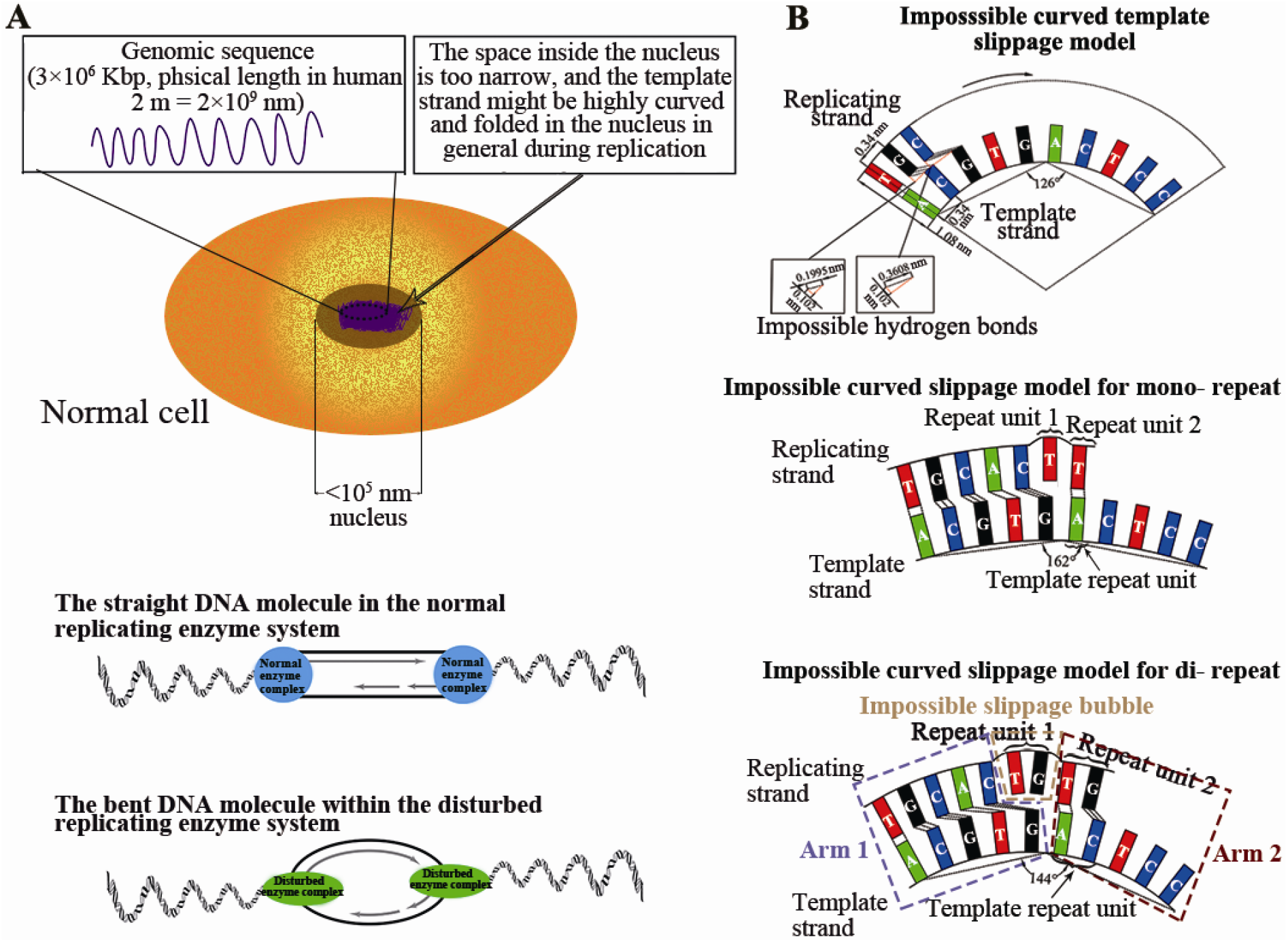
The DNA molecule is highly curved or folded in the nucleus and the impossible curved slippage model. A. Schematic diagram of the size of the nuclear space (top) (Lamond, 2002); The normal replicating enzymes complex straighten the DNA molecule, while the disturbed replicating enzymes complex may cause the DNA molecule return to curved state (bottom). B. Impossible curved template slippage model according to the strict geometric calculation of the space of a nucleotide and the stability of hydrogen and phosphodiester bonds (top); Mono- and dinucleotide repeats may be impossibly produced in curved replicating strands (middle and bottom).

Firstly, we proposed a curved template slippage model. When the curved DNA strand is used as the template strand on inner side, the replication strand is longer than the template strand and can form more nucleotides than the template strand on the outside for during replication process. The replication strand should be longer than template strand, then, is able to provide extra spaces for accommodating the extra repeat bases (Fig 3B). However, it is well known that the links of base pairs mainly depend on 2 types of hydrogen bonds, N—H…:N and N—H…:O (Heyrovska, 2006), and the strengths of these hydrogen bonds are negatively correlated to the distance between every base pair; the strength of the hydrogen bond is about 3% of the 3’, 5’-phosphodiester bonds (Gao et al, 2004; Griffiths et al, 2000; Luo, 2007; Wang, 1993) (Fig 2A), so the distance between the bases is fixed; even if there is space to form a slippage bubble, the hydrogen bond should be elongated to exceed the threshold of 0.167 nm (Heyrovska, 2006) and should be easy to be broken off in such condition. So, the curved slippage model is able to provide spaces for forming slippage bubble with forming unstable hydrogen bonds double-chain structures (Arm1 and Arm2) at both sides of the slippage bubble (Fig. 3B, Fig S2), indicating that the curved slippage model should be unreasonable.

Then we proposed a folded slippage model. In this case, the folded template strand forms a slippage bubble above the folding site to have sufficient space for accommodating the repeat nucleotides in replication process, the phosphodiester bonds are not elongated, but the bases are well paired with the stable hydrogen bonds at both sides of the slippage bubble (Fig 4). If folding angle is proper, thereby it is most possibly to form a very stable double-stranded folded slippage structure to provide chances for producing repeats, with considering nucleotide geometric spaces and stability of phosphodiester and hydrogen bonds. Actually, there are two conditions of the folded slippage models: When template strand is on the inner side, the repeat unit duplicated to produce new repetitive unit or repeat expansion (Fig 4); and when the template strand is on the outside, the replication strand may make the repetitive sequences to contract (Fig 5); the features of this folded slippage model can easily explain the widely observed microsatellite mutations with expansion and contraction of repeat units (Ellegren, 2004; Gemayel et al, 2010; Gymrek et al, 2016; Kelkar et al, 2011; Mirkin, 2007). In addition, replication slippage of template strands with different folding angles may result in the expansion or contraction of repeat units with different sizes. When template chains are folded on the inner side at a rotation angel of 18°, 36°, 54°, 72°, 90° and 108°, the replication strands will produce mononucleotide to hexanucleotide repeat expanding respectively (Fig 4). So, it is necessary to break off the number of hydrogen bonds from 2 to 18 without elongating the phosphodiester bond to produce repeats; it suggested that the difficulty of formation repeats from mono- to hexanucleotide is gradually increasing, and also means the occurrence of mono-, di-, tri-, tetra-, penta- and hexanucleotide repeat is gradually decreasing; that is well consistent with our statistic data (Table 1, Table S2). Vice versa, when template chains are folded on the outside at a rotation angel of 18°, 36°, 54°, 72°, 90° and 108°, the replication strands will produce responding repeats contracting respectively (Fig 5). These features are well corresponding to the microsatellites which usually refers to the tandem repeats with repeat units from mono- to hexanucleotides(Ellegren, 2004; Kelkar et al, 2010; Zhao et al, 2011). According to this rule, we also describe the possible folded template slippage models of hepta-, otca-, nona- and decanucleotide repeats (Figs S3 and S4). In fact, the replicating strand must break off at least from 14 to 30 hydrogen bonds to make a folded slippage bubble, the energy to break off so much hydrogen bonds are almost close to energy of phosphodiester bond, then, they are very difficult to occur, and therefore, this is consistent with the observations that such long tandem repetitive sequences are often not very abundant in the genomes (Gemayel et al, 2010; Legendre et al, 2007). The (A_m_T_n_) repeats growing faster than (G_m_C_n_) repeats also suggested that the broken number of hydrogen bonds involves in the speed of repeat expansion (Katti et al, 2001; Schlötterer & Tautz, 1992; Sinai et al, 2019; Tian et al, 2011). Although this folded slippage model is just simply described in a plane form, it can still clearly simulate and explain the repeat sequences producing process. We also use the same space size to make the double-helical three-dimensional forms show the folded slippage model more intuitively (Figs 4 and 5), and the precise folding angle in the three-dimensionally double-helical forms and other issues desire further study.

**Figure 4.**
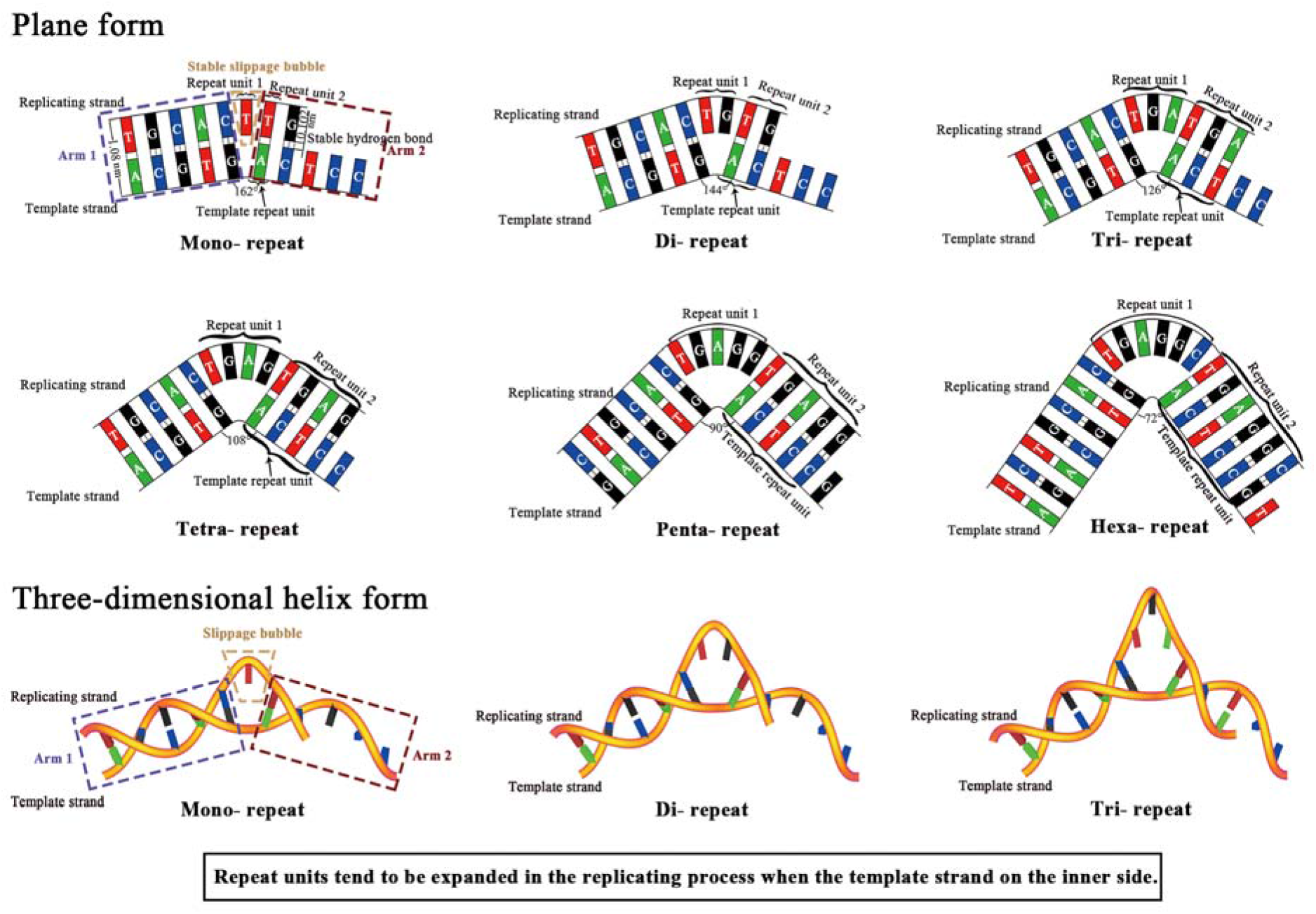
Stable folded slippage models of mononucleotide to hexanucleotide repeats amplification according to the strict geometric calculation of the space of a nucleotide and the stability of hydrogen and phosphodiester bonds. Repeat units tend to be expanded in the replicating strands when the template strands are on the inner side of the folded slippage models respectively. The bottom 3 figures were the folded slippage models in three-dimensional helix form.

**Figure 5.**
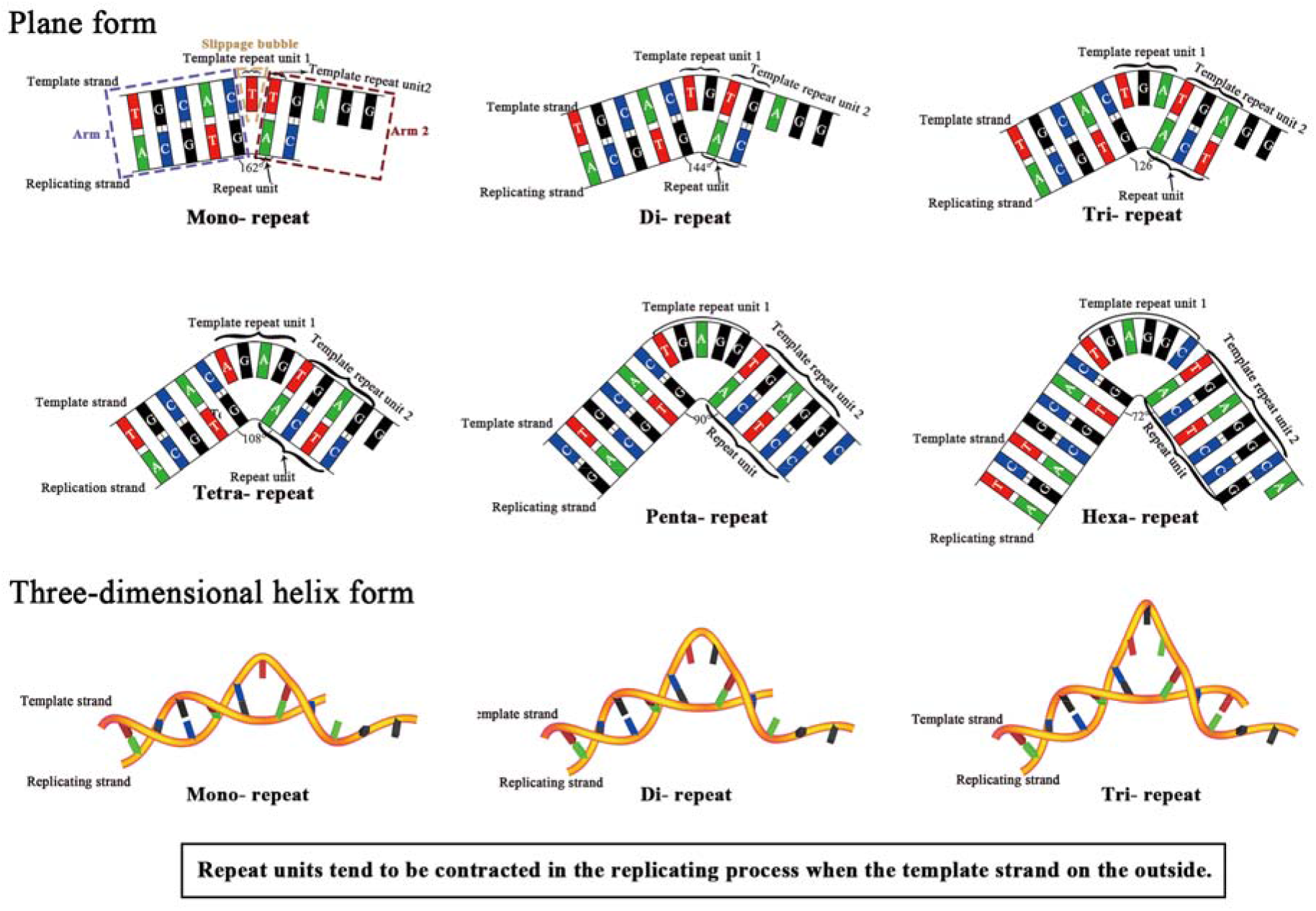
Stable folded slippage models of mononucleotide to hexanucleotide repeats contraction according to the strict geometric calculation of the space of a nucleotide and the stability of hydrogen and phosphodiester bonds. Repeat units tend to be subtracted in the replicating strands when the template strands are on the outside of the folded slippage models respectively. The bottom 3 figures were the folded slippage models in three-dimensional helix form.

There is enough geometric space in the slippage bubble of the folded template model to accommodate repeat nucleotides without stretching the phosphodiester bonds, compared with the straight template slippage model. In contrast to the curved template model, the difference in the folded model is that the two sides of the slippage bubble are stably paired, and the Arm1 and Arm2 similar to the straight template replication model are formed at both sides (Figs 4 and 5). The folded model takes full account of the space required by nucleotides, the stability of phosphodiester bonds and the strength comparison between phosphodiester bonds and hydrogen bond, and is easy used to explain microsatellite mutations with repeat unit expansion and contraction. Therefore, we propose that the folded template chain slippage model may be considered as the most reasonable model for explaining repeats production in replicating process, and the folded template strand slippage model may be responsible for the continual producing of repeat sequences and the remaining of high percentage of repeat sequences in genomes.

### Microsatellites tend to expand

As stated above, according to the folded slippage model, template chain folding on the inner side may make the replicating chain slippage for repeats expansion, vice versa, the template chain folding on the outside may make the replicating chain slippage for repeats contraction; and it seems that the possibility of repeats expansion and contraction is same. However, there are two manners for the repeat sequences contraction, one is above mentioned the template chain folds on outside, another is also above stated general mutations; the high content of the repeat sequence is still in a stable state in the genome of each species, suggesting that the possibility of repeat expansion should be higher than repeat contraction. And many reports also suggest that there is a higher possibility of repeat expansion than repeat contraction (Fungtammasan et al, 2015; Fungtammasan et al, 2016; Neil et al, 2018).

When the folded template chain slippage was deeply investigated, the replicating straight template DNA chain should return to folded under external forces from the narrow and crowded cell nucleus when the replicating enzyme complexes are disturbed, and usually the replicating enzyme complexes may provide power for balancing the external forces to drag the template DNA molecule straight. Then, we proposed an external force function for template strand returning to folded, and this function may be helpful to explore the probability of expansion and contraction. When the template strand is on the inner side, the nucleotide bases are outward, and the space of bases at the folded site become wide and loose at outward part; while it is on the outside, the base in the folding position is squeezed inward. Comprehensively considering the small difference of the space of nucleotides at the folded site, it can be easy accepted that the external forces to make template strand folded with bases loose should be smaller than that to be squeezed; therefore, the external force required for the template strand folded on the outside (F^o^) is inevitable greater than that (F^i^) on the inner side, it can be described as F^o^ >F^i^, suggesting that the probability for the template strand folded on the inner side is higher than that on the outside; as our folded slippage model suggested that the repeats tend to expand when the template strand on inner side and contract when the template strand on outside, therefore, the possibility of repeat expansion (P^e^) is most possibly higher than that for repeat contraction (P^c^), it can be described as P^e^>P^c^ (Fig 6). The SSR studies, like in Huntington disease related locus and myotonic dystrophy type 1 locus, all showed SSR expansion biased (Higham et al, 2012; Larson et al, 2015; Macdonald et al, 1993; Mirkin, 2007; Sznajder & Swanson, 2019), which proving that the expansion and of short SSRs are more frequent than that of contraction.

**Figure 6.**
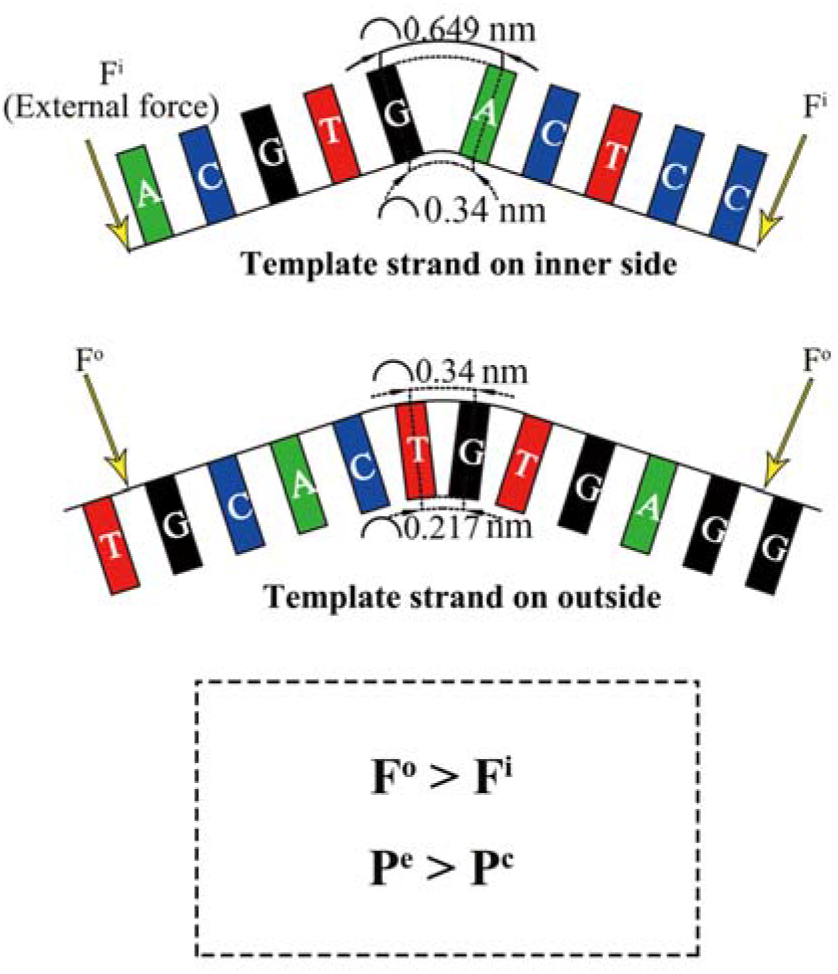
Repeats production incline to expansion. F^o^, F^i^ refer to the force required for the two template strands to bend, respectively. F^o^>F^i^ means that the force of the template strand bending downward is greater than the bending upward, and P^e^>P^c^ means that the possibility of the template strand bending upward is greater than the downward bending.

Thus, according to formula (2):

When the template strand on the outside, repeats tend to contract, so *λ^c^* < 0, thus, 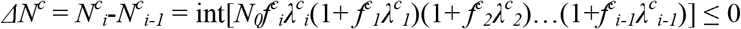.

When the template strand on the inner side, repeats tend to expand, so *λ^e^* > 0, thus, 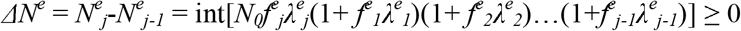.

The general repeat expansion and contraction can be described as:

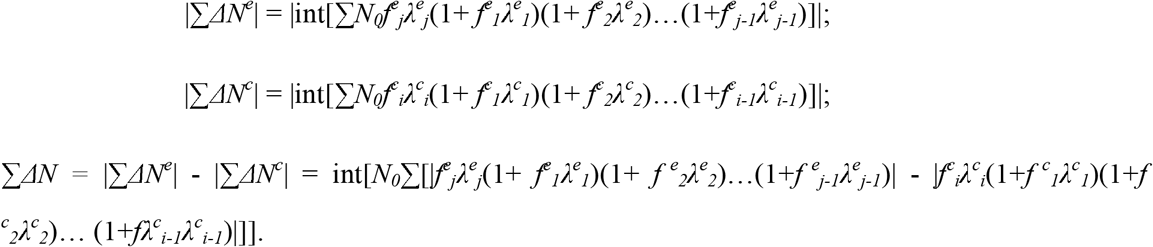

Because *λ* was defined as coefficient of occurring repeats, the possibility of repeat expansion (P^e^) is positively proportional to *λ^e^* and the possibility of contraction (P^c^) is positively proportional to the absolute value *λ^c^* (|*λ^c^*|), if we suppose that *f^e^ = f^e^ = f, i = j*, and as generally P^e^ > P^c^, then *λ^e^* > |*λ^c^*|, and also 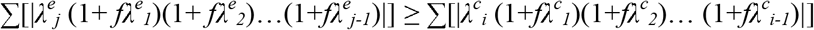, therefore, 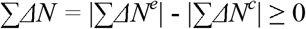.

So, when the external forces for returning the folded template strand were considered, the possibility of repeat expansion should be higher than that of repeat contraction, then the revised formula (2) is also able to explain the remaining of high percentage of short repeats in genomes under a mechanism of continually producing repeats; and this mechanism might result from the folded template chain slippage model, which is possibly responsible for the widely occurring short tandem repeats, also called microsatellites or SSRs in eukaryotic, prokaryotic and also viral genomes. We improved the straight slippage model to folded slippage model by fully considering the geometric spaces of nucleotides base, the relationship between phosphodiester and hydrogen bond and the stability of these bonds. The slippage model showed that the straight replicating template DNA may return to be some extent of folded resulting from disturbed replicating enzyme complexes, and may provide chances for continually producing much amount of short repeats; though the long unit repeats may be related with the former slippage model (Gemayel et al, 2010; Viguera et al, 2001). The easily forming of folded slippage may be also responsible for the widely observed fact that repetitive part of genome is usually evolved hundred or more times than other part with only repeat units expansion and contraction (Giesselmann et al, 2019; Kelkar et al, 2011; Kim et al, 2013; Mandal et al, 2019), though the repeats occurred more in non-coding regions than in coding regions possibly because of different selective pressures (Ellegren, 2004; Gemayel et al, 2010; Mirkin, 2007). Most of new occurring repeats should be lethal mutation and may have been negatively selected to lost; some of new occurring repeats should be deleterious in genomes and responsible for a series of diseases (Arturo et al, 2010; Larson et al, 2015; Sun et al, 2018; Sznajder & Swanson, 2019); many neutral repeat expansions may be lost or fixed with no functions in genomes by genetic drift (Muller et al, 2014); and some beneficial repeat expansions may promote the emergence of different new properties or functions, that is why the repeat sequences are reported with so many different roles (Gymrek et al, 2016; Hannan, 2018; Hood et al, 1996; Li et al, 2004; Mrazek, 2006; Sinai et al, 2019; Vinces et al, 2009b). And the longer repeats might originate from short repeat expansion by the folded template slippage, and the longer genomes possibly evolved from the short genome with related to the continuous repeats producing folded slippage model in the long evolutionary replicating process.

## Materials and Methods

### Sequences resource

We downloaded 55 genomic sequences from Genbank of a fairly wide range of species that covering animals, plants, fungi, protozoa, bacteria, archaea and viruses. The segments for SSR analysis were randomly selected from different regions of these 55 genomic sequences, which range from 3000 to 96600 bp in length and do not contain any gaps, to verify the widespread distributions of SSRs, as the full genomic sequences are too long.

### Repeat extraction

The perfect simple sequence repeats were extracted by Imperfect Microsatellite Extraction Webserver (IMEx-web, http://imex.cdfd.org.in/IMEX/index.html) from those 55 randomly-selected reported segments. The minimum iterations for all perfect mono- to hexanucleotide repeats were set at 3, 2, 2, 2, 2, 2 to mine the data more completely in this study, comparing with most researchers setting iterations at relatively higher self-defined values, and 3 iterations for mononucleotide repeats were confined to ensure to be commonly recognized as the SSRs.

### Null hypothesis test

We also extracted perfect mono- to hexanucleotide repeats under the above threshold in the sequences that were generated by a program written in C language (Program S1) according to the nucleotide compositions and sizes of those 55 reported segments. Then, the validating test, which can verify that the short SSRs extracted in those 55 reported segments are nonrandom sequences, was based on the comparison of the SSR percentages in the reported segments and our generated segments.

### Model drawing of DNA replication

Different models were drawn to simulate the DNA replication. Normally in straight model, the hydrogen bond length between 2 paired nucleotides is reported to be 0.102 nm and the distance between 2 neighboring nucleotides is 0.34 nm, importantly, owing to the nucleotides occupying almost same space in DNA strands, the space of a nucleotide was simplified into a geometric plane form in this analysis, which was 0.489 nm in length and 0.34 nm in width. Then we applied AutoCAD to draw the straight, curved and folded slippage models according to the strict geometric calculation of the spaces of nucleotides and different strengths between hydrogen bonds and phosphodiester bonds. And the slippage models in helix structure were achieved by Rhino, which is an industrial drawing software.

## Availability of data and materials

Supplementary Tables are online at https://github.com/DooYal/Supplementary-Table-for-submitting-relatively-...-/tree/DooYal-patch-manuscript_folded/supplementary%20tables

## Acknowledgments

The authors thank Qijun Tian, who is from School of design in Hunan University, for his technological help at designing the folded slippage model in helix structure.

## Author contributions

Z. Tan designed and directed this study. D. Li, S. Pan and H. Zhang performed the data analysis, Y. Fu, Z. Peng, H. Zhang and L. Zhang helped for performing data analysis. F. Xu helped for mathematic calculation. S. Peng, Hanrou Huang and Ruixue Shi helped for drawing maps. Z. Tan and D. Li prepared the manuscript. H. Zhang and S. Peng helped for the manuscript preparation.

Supplementary Table are online at https://github.com/DooYal/Supplementary-Table-for-submitting-relatively-...-/tree/DooYal-patch-manuscript_folded/supplementary%20tables

## Supplementary Figures

**Figure S1.**
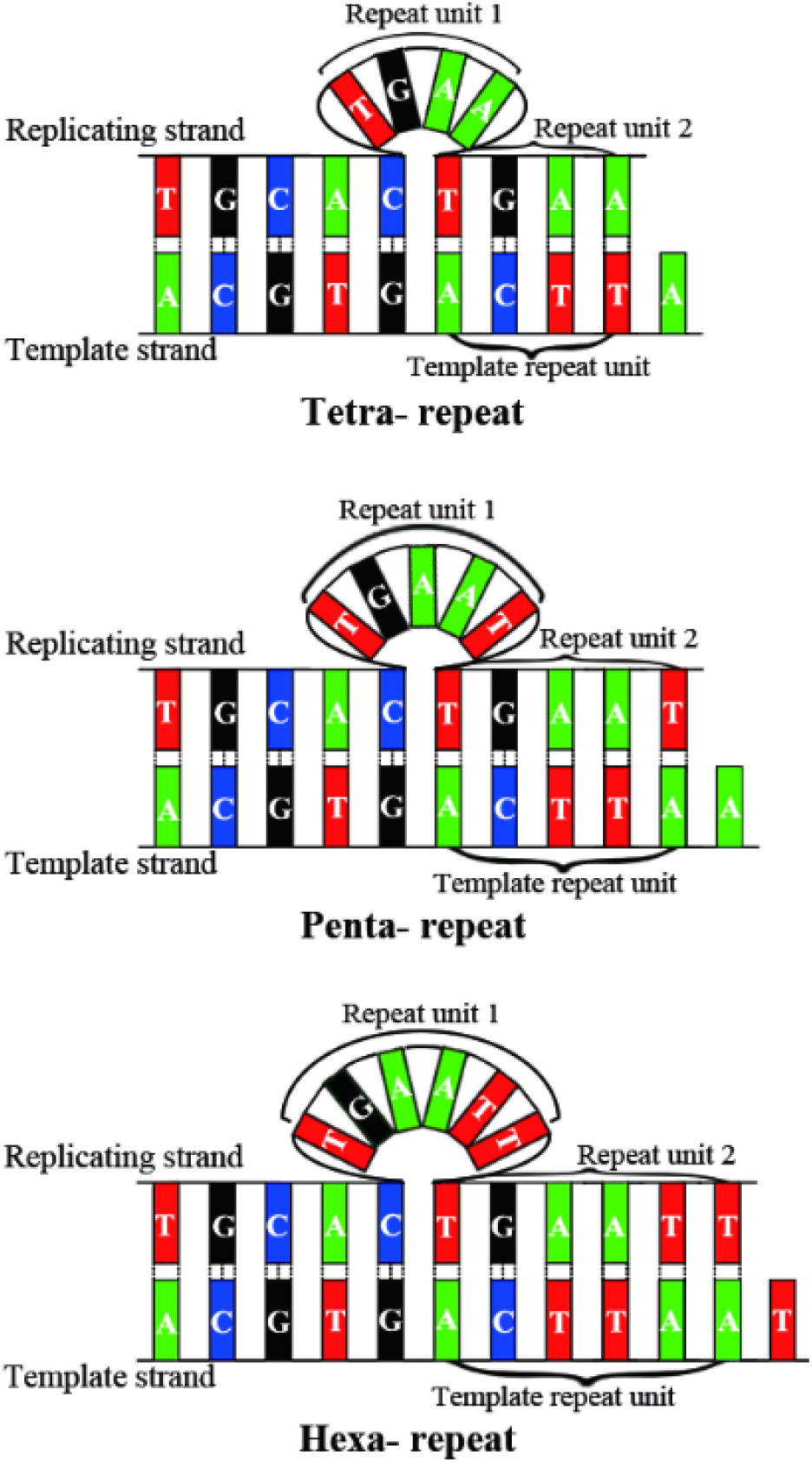
Impossible straight slippage models for tetra- to hexanucleotide repeats when the slippage bubble occurs at the replicating strand. The model drawing was based on the strict geometric calculation of the space of a nucleotide and the stability of hydrogen and phosphodiester bonds.

**Figure S2.**
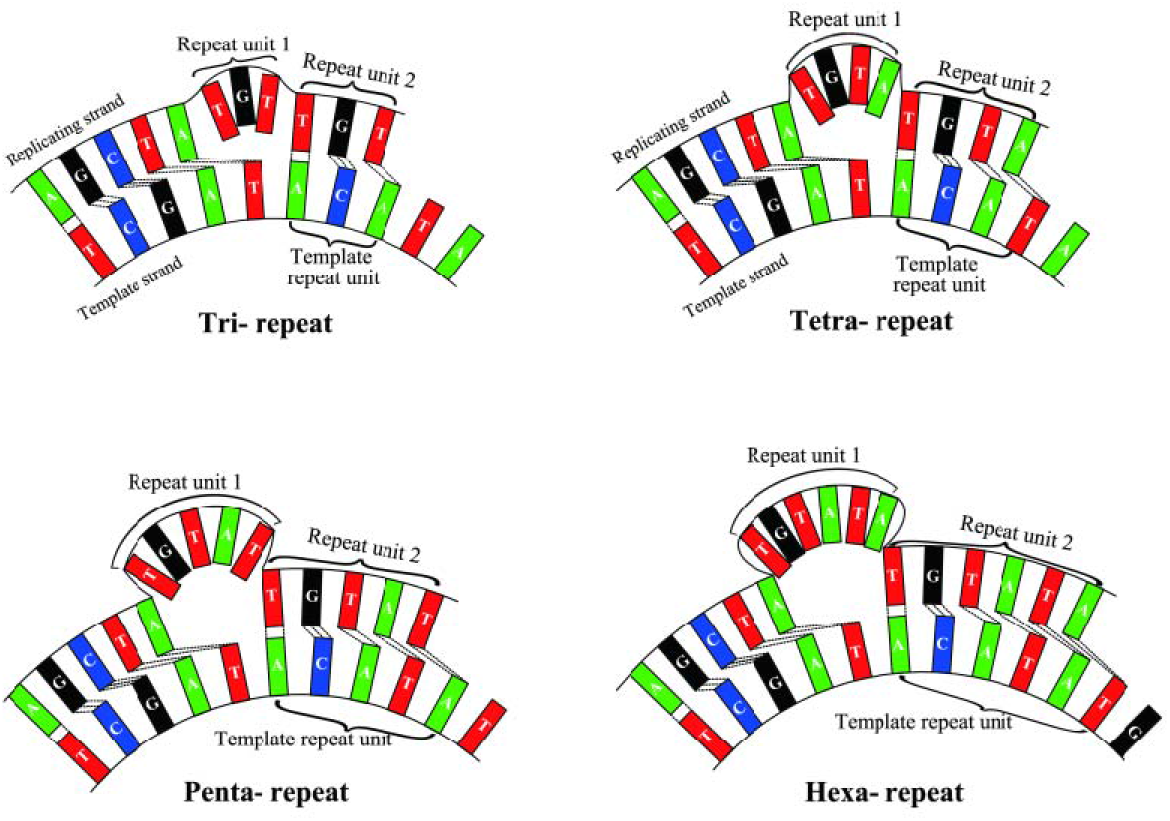
Impossible curved slippage models for tri- to hexanucleotide repeats when the template strand in the inner side of the models.

**Figure S3.**
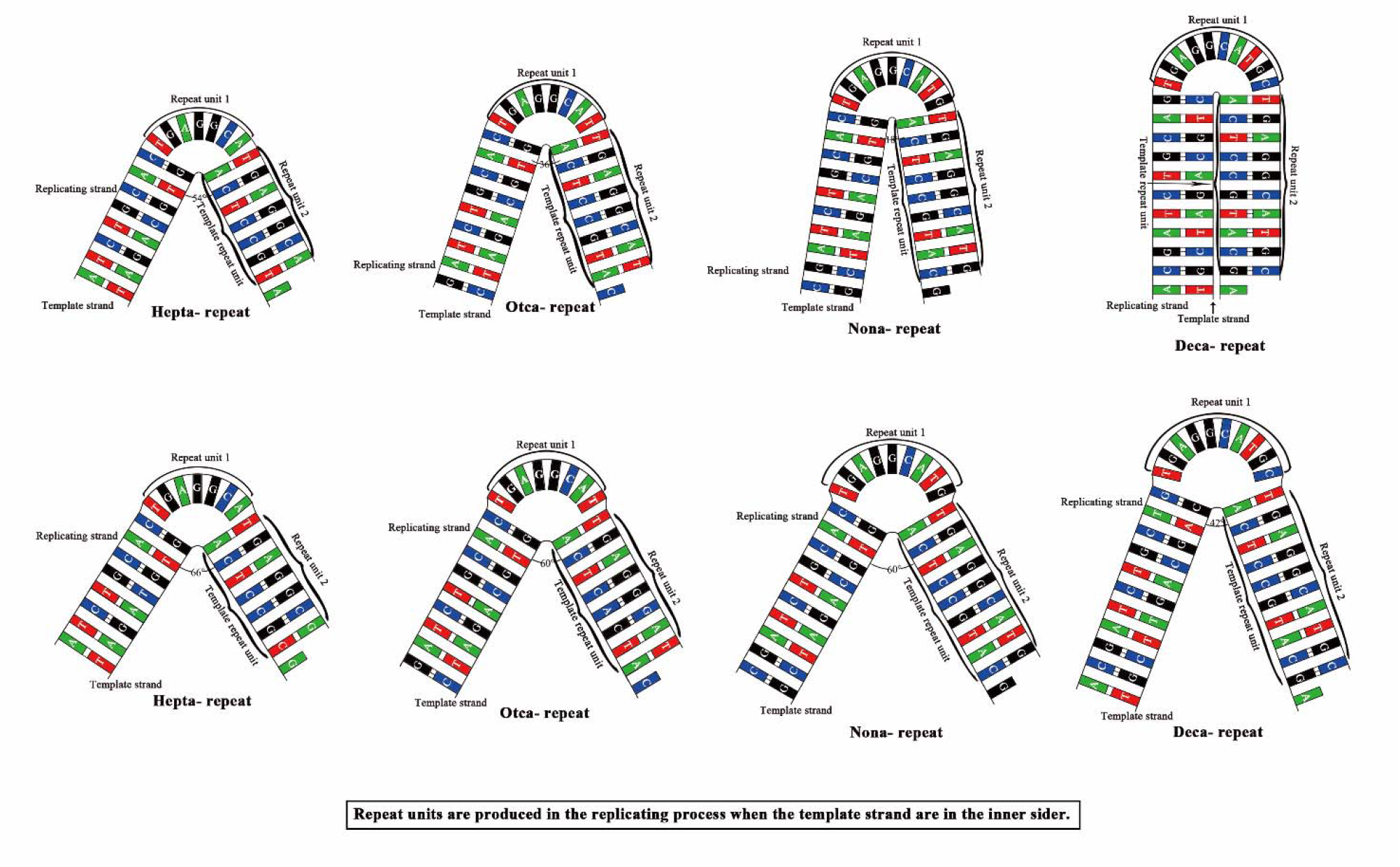
The possible folded slippage models for hepta- to decanucleotide repeat amplification. Repeat units tend to be expanded in the replicating strands when the template strands are on the inner side of the folded slippage models respectively.

**Figure S4.**
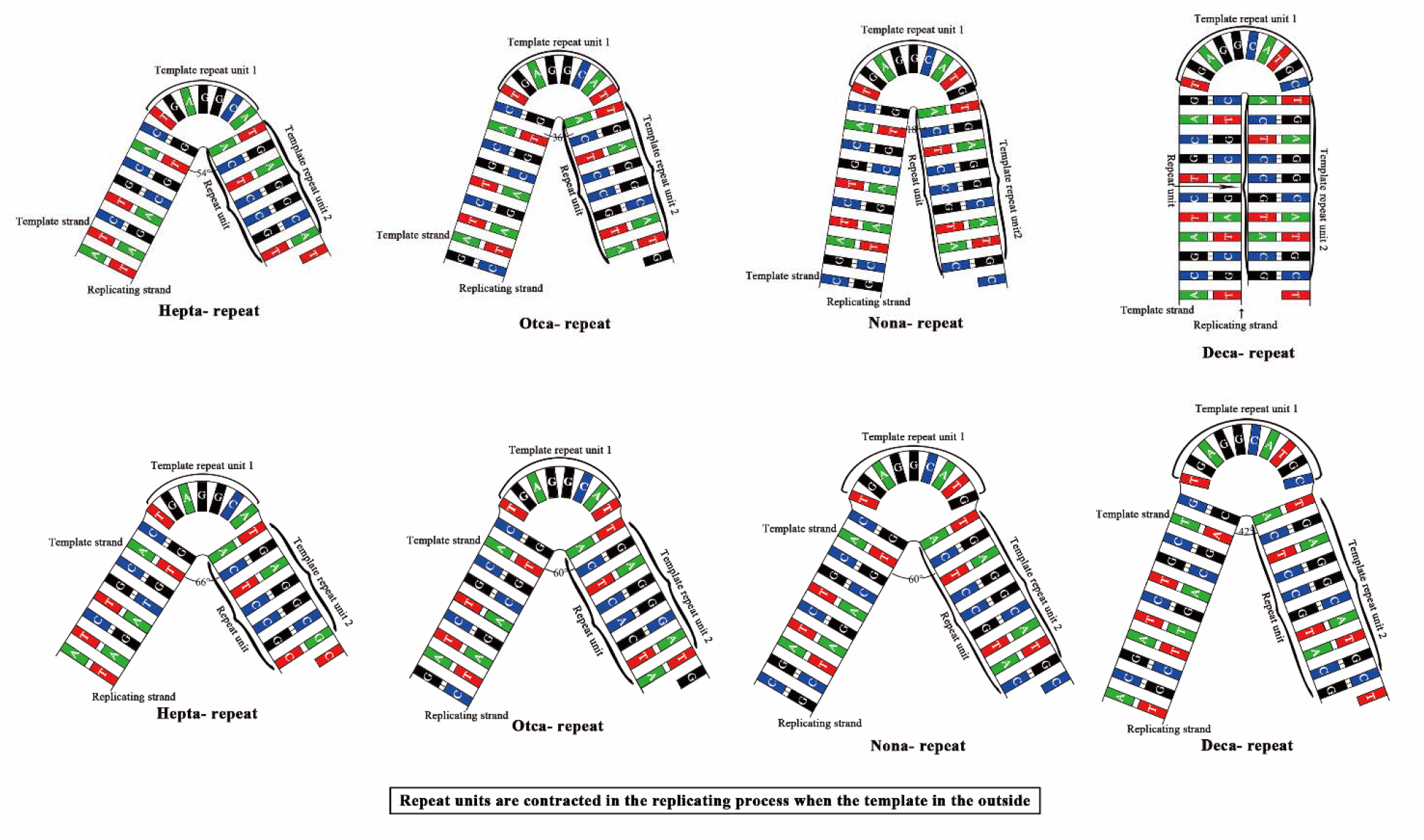
The possible folded slippage models for hepta- to decanucleotide repeat contraction. Repeat units tend to be subtracted in the replicating strands when the template strands are on the outside of the folded slippage models respectively.

